# Dissociable neural signals for reward and emotion prediction errors

**DOI:** 10.1101/2024.01.24.577042

**Authors:** Joseph Heffner, Romy Frömer, Matthew R. Nassar, Oriel FeldmanHall

## Abstract

Reinforcement learning models focus on reward prediction errors (PEs) as the driver of behavior. However, recent evidence indicates that deviations from emotion expectations, termed affective PEs, play a crucial role in shaping behavior. Whether there is neural separability between emotion and reward signals remains unknown. We employ electroencephalography during social learning to investigate the neural signatures of reward and affective PEs. Behavioral results reveal that while affective PEs predict choices when little is known about how a partner will behave, reward PEs become more predictive overtime as uncertainty about a partner’s behavior diminishes. This functional dissociation is mirrored neurally by engagement of distinct event-related potentials. The FRN indexes reward PEs while the P3b tracks affective PEs. Only the P3b predicts subsequent choices, highlighting the mechanistic influence of affective PEs during social learning. These findings present evidence for a neurobiologically viable emotion learning signal that is distinguishable—behaviorally and neurally—from reward.

**Significance:** For nearly a century, scientists have asked how humans learn about their worlds. Learning models borrowed from computer science—namely, reinforcement learning—provide an elegant and simple framework that showcases how reward prediction errors are used to update one’s knowledge about the environment. However, a fundamental question persists: what exactly is ‘reward’? This gap in knowledge is problematic, especially when we consider the multiplicity of social contexts where external rewards must be contextualized to gain value and meaning. We leverage electroencephalography to interrogate the role of emotion prediction errors—violations of emotional expectations—during learning. We observe distinct neural signals for reward and emotion prediction errors, suggesting that emotions may act as a bridge between external rewards and subjective value.

## Introduction

Reward prediction errors (PE)—the difference between expected and experienced reward—serve as the dominant mechanism for explaining how people learn to make adaptive, value-based decisions^1–6^. Reward PEs function within a reinforcement learning (RL) framework, illustrating how people adjust their actions based on past experiences to achieve more successful outcomes^7^. This framework has been successfully applied to explain a host of simple behaviors such as avoiding financial losses^1^ and navigating new environments^8^, to more complex behaviors, such as determining who can be trusted^9^. While the term reward is frequently used to explain learning, the fact that ‘rewards’ in the social world are abstract, difficult to quantify, and shaped by multiple features of a social situation, suggests a gap between the externally ‘rewarding’ reinforcers encountered (e.g., money, smiles) and how the brain interprets them as value. A critical unresolved question thus centers around what exactly is ‘reward’ and how does the brain represent it? Here we examine how the human brain computes external rewards into internal value during a social learning paradigm.

An intuitive possibility for how external reinforcers are transformed into internal value comes from the field of emotion. Decades of work indicate that emotions play a vital role in the decision-making process^10,11^, where stress inductions^12^, mood inductions^13^ and emotion regulation^14^ can all impact choice. Indeed, several affective theories propose that emotions *are* the evaluation of external rewards, and thus possess the capacity to influence future behavior^15–18^. Building on this theory, our previous research operationalized affect within an RL framework, which led to the hypothesis that violations of emotion expectations—known as affective prediction errors (PEs)— influence choice. By formally quantifying affective PEs as the difference between expected and experienced emotion, we observed that affective PEs exhibit an independent effect that is stronger than monetary reward PEs in predicting one-shot social choices^19^. The distinction between emotion and reward was further exhibited in an independent sample, where individuals at risk of depression demonstrated selective impaired use of affective PEs but fully intact use of reward PEs in a social exchange task.

Although this dissociation suggests affective PEs have a central role in guiding socially adaptive behaviors, it remains unknown whether affective PEs also act as critical signals during trial-by-trial learning, where knowledge about others must be continually updated to adjust future choices. Additionally, there is no evidence for the separability between affective and reward PEs at the neural level. Neural separability—at the temporal, functional, or localization levels—between affective and reward PEs would be strong evidence that emotion serves as a distinct learning signal separate from reward, one that may transform external rewards into internal, subjective value.

To test whether affective and reward PEs are critical for learning and are separable at the neural level, we used a repeated social exchange paradigm in conjunction with electroencephalography (EEG). EEG was chosen for its superior temporal resolution capable of capturing unfolding neural processes on the order of milliseconds. We *a priori* identified three potential event-related potentials (ERPs) known to reflect changes in EEG activity in response to feedback or reward processing. In particular, the feedback-related negativity (FRN) is an ERP thought to reflect the evaluation of surprising events^20,21^ and is theorized to be the neural basis of reward PE processing^22^. Additionally, the P3a and P3b are commonly linked to various aspects of feedback processing, including reward magnitude^23^, reward valence^24^, and rare, surprising outcomes^25^. Given the mixed evidence of how these ERPs map onto the construct of reward, we were agnostic as to which neural signal would preferentially index reward or affective PEs.

EEG was recorded during a repeated Ultimatum Game (UG), where participants (N=41) interacted with three different partner types offering a range of fair to unfair monetary offers (fair, unfair, neutral; see Methods and Fig. 1B). The repeated nature of the UG, five trials in a row per partner, allowed participants to update their expectations of a partner based on the history of offers with that person. We included two key measurements to enhance our understanding of how rewards and emotions influence feedback processing and updating (Fig. 1A). First, rather than using computational models to infer participants’ reward expectations^26,27^, we asked them to report the amount of money they expected to receive on each trial (ranging from $0 to $10). This allowed us to compute trial-by-trial reward PEs as the discrepancy between the actual offer and the expected one. Second, participants used a 2D affect grid to predict how they thought they would feel after receiving an offer (affect expectation), and to express how they actually feel once the offer was received (affective experience)^19,28^. This measure captures participant’s core affect, a consciously accessible facet of subjective feelings that categorizes feelings into core dimensions of valence (pleasurableness) and arousal (alertness/intensity). We computed affective PEs for both arousal and valence dimensions as the difference in participant’s expectation and experience on a trial-by-trial basis (Fig. 1C). Together, these measurements allow us to map all three empirical PEs (reward, valence, and arousal) onto distinct neural EEG signatures expressed during social interactions (Fig. 1D).

**Figure 1.**
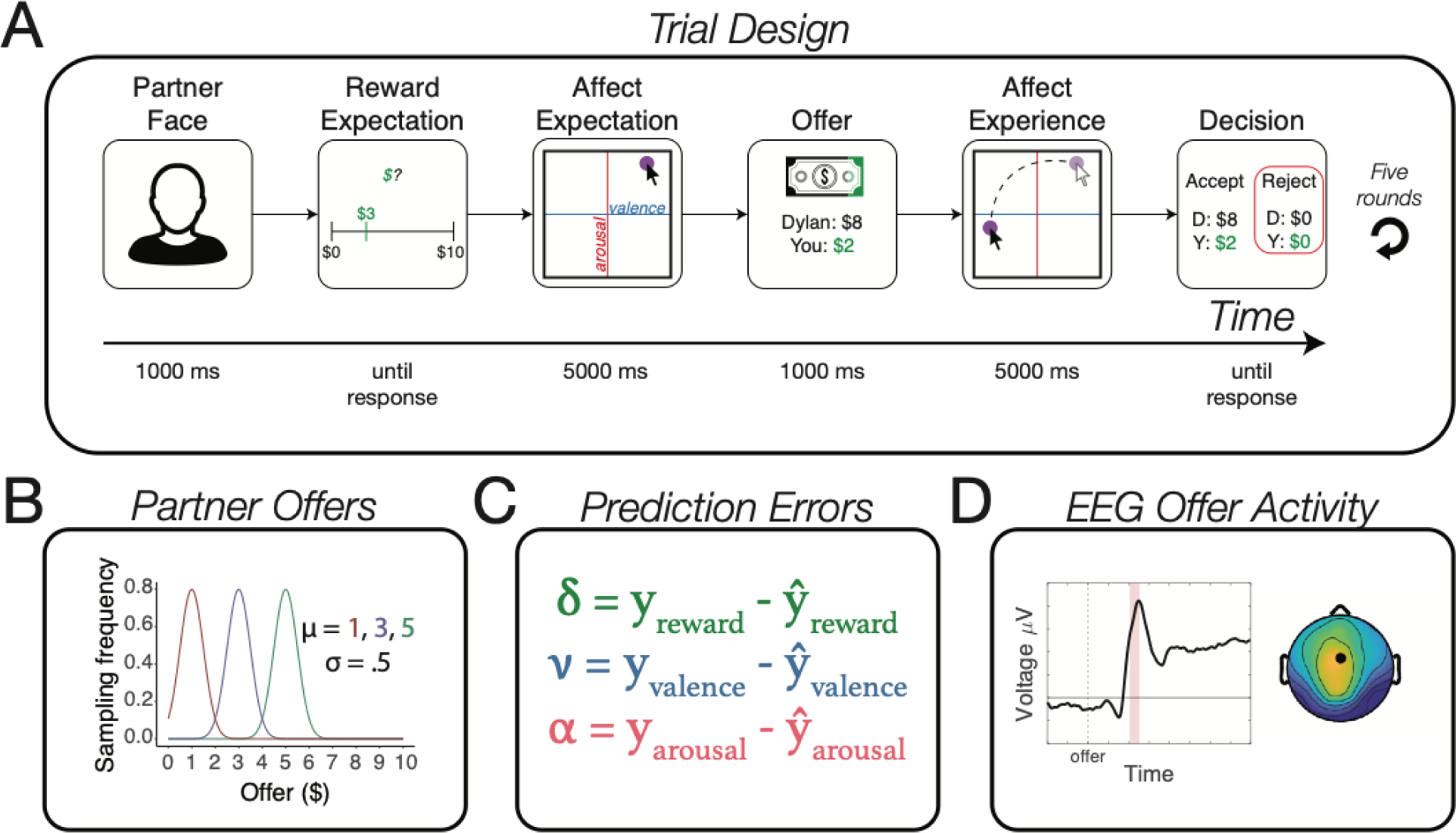
Repeated Ultimatum Game (UG) Design. **A) Trial Design.** Participants are partnered with another individual and play five rounds of a repeated UG. Participants first see a photograph and name of their partner (Partner Face), before being asked to report how much money they expected to receive in an offer (Reward Expectation) and how they expected to feel after the offer (Affect Expectation). Next, participants received an offer showing the proposed amount for the participant as well as the amount kept by the partner (Offer). After receiving the offer, participants indicated how they felt (Affective Experience) before deciding to accept or reject the offer (Decision). Timings show the average maximum duration of each of the stages, with “until response” meaning the program waited until the participant gave a response. **B) Partner Offers.** Unknown to participants, offers were determined by different normal distributions per partner type. On average, unfair partners gave $1, neutral gave $3, and fair gave $5; all partners’ normal distributions used a standard deviation of $0.50. **C) Prediction Errors (PEs).** On each trial we compute three empirical PEs: a reward PE (δ), a valence PE (ν) and an arousal PE (α). In the equations, ŷ refers to an individual’s prediction about the reward or emotion they would experience, while y refers to their actual experience. **D) EEG Offer Activity.** Our analyses focus on the EEG activity occurring after the offer presentation. The average EEG activity at electrode FCz in response to offers is shown alongside a topography of the average activity across electrodes between 200ms to 250ms.

## Results

### Reward and affective PEs have separable contributions to learning

Given our prior work found that valence, compared to reward PEs, exert a stronger influence on one-off decisions to punish norm violators^19^, we began by examining the marginal strength of reward, valence and arousal PE signals when deciding to punish in a social learning context. This additive linear mixed-effects model (LMM) represents a strict test of our theory since each PE type competes with all others to explain variance during learning. Replicating our prior research, we found that both valence (β = −0.76 ± 0.11, *z* = −6.65, *P* < 0.001) and reward PEs (β = −0.67 ± 0.14, *z* = −4.75, *P* < 0.001) have independent contributions when learning when to punish. That is, participants punished at higher rates when experiencing more unpleasantness (valence) or less reward than expected. Unlike our prior results, however, we observed no unique contribution of arousal PEs on decisions to punish (β = 0.13 ± 0.11, *z* = 1.14, *P* = 0.26) once valence and reward PEs are accounted for.

To examine how the relationship between each PE type and choice changes over time, we interacted PE type with round number. We observe that the strength of reward and valence PEs change in opposite directions overtime (Table 1; Fig. 2A). Valence PEs exert the strongest relationship to choice on the first round when uncertainty is greatest, and significantly weakens over time as uncertainty about a partner’s behavior is slowly resolved. In contrast, reward PEs show a marginally significant interaction with round number such that they weakly predict choice in the beginning but become more predictive over time. Directly pitting both PE types against one another reveals that valence PEs have a significantly stronger impact on motivating punitive choices on the first round when compared to reward PEs (β coefficient test: *z* = -1.70, *P* = 0.04), while reward PEs have a stronger, albeit not-significant, influence on the final round, when compared to valence PEs (*z* = 1.05, *P* = 0.15). This reversal reveals how valence PEs are more impactful early on when uncertainty is greatest. Once participants have better estimates about the type of offer to expect, reward PEs become more useful for informing whether to accept or reject an unfair offer.

**Table 1.** Separable effects of valence and reward PEs predict learning to punish.

| Variable | Estimate (SE) | z | p |
| --- | --- | --- | --- |
| Punish |  |  |  |
| Intercept | -1.61 (0.32) | -5.03 | <.001*** |
| Reward PE | -0.50 (0.17) | -3.02 | .003** |
| Valence PE | -0.99 (0.15) | -6.42 | <.001*** |
| Arousal PE | 0.05 (0.14) | 0.34 | .737 |
| Round | 0.04 (0.03) | 1.66 | .10 |
| Reward PE×Round | -0.06 (0.03) | -1.87 | .06 |
| Valence PE×Round | 0.08 (0.03) | 2.30 | .02* |
| Arousal PE×Round | 0.03 (0.03) | 0.98 | .33 |

*Note.* Reward PEs are calculated by taking the difference between the experienced and predicted reward. Valence PEs and Arousal PEs are calculated by taking the difference between the experienced and predicted emotion on the relevant affect dimension. All variables were scaled but not mean-centered, as the zero point on each scale refers to the meaningful instance where expectations matched experience. The model includes subject-specific random intercepts and slopes for Reward PE, Valence PE, and Arousal PE. The dataset includes 7,376 observations from 41 participants. \*\*\* $p < .001$ , \*\* $p < .01$ , \* $p < .05$ .

**Figure 2.**
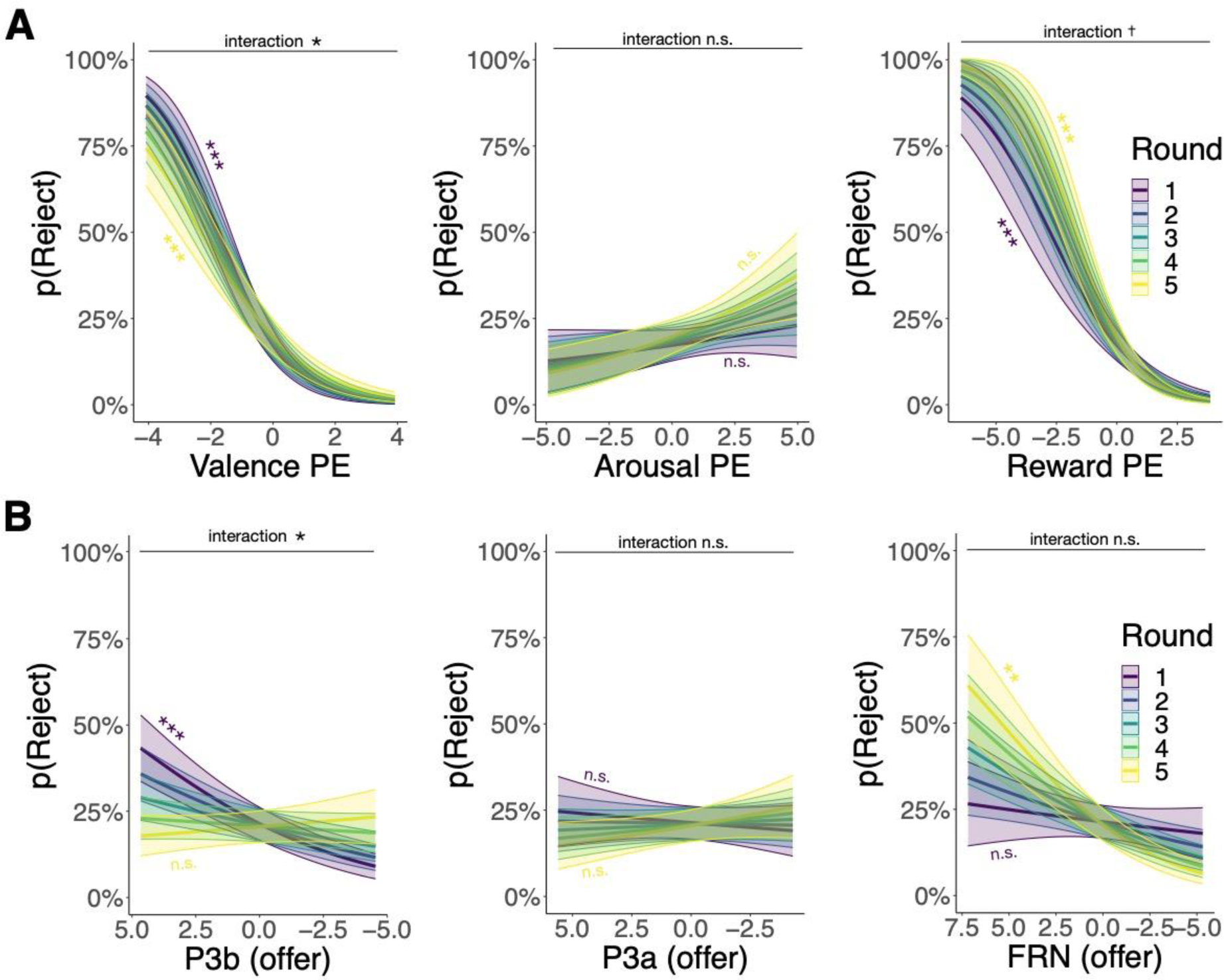
Separable behavioral and neural effects of valence and reward PEs predict learning to punish. **A) Valence and reward PEs predict choice differently across rounds.** The data on each graph reflect the probability of rejecting the offer from Table 1 and the colour of each line indicates the round number (1 – 5) of the Ultimatum Game. Negative values reflect negative PEs, indicating less pleasantness (valence), arousal, and money (reward) than expected. **B) Relationship between ERP and choice over round.** The data on each graph reflect the probability of rejecting the offer from Table 2 and the colour of each line indicates the round number (1 – 5) of the Ultimatum Game. Positive values reflect positive EEG amplitudes, indicating a greater P3b, P3a or FRN effect. Shaded areas reflect ±1 S.E. ***P<.001, **P<.01, *P<.05, t = 0.06.

**Table 2.**
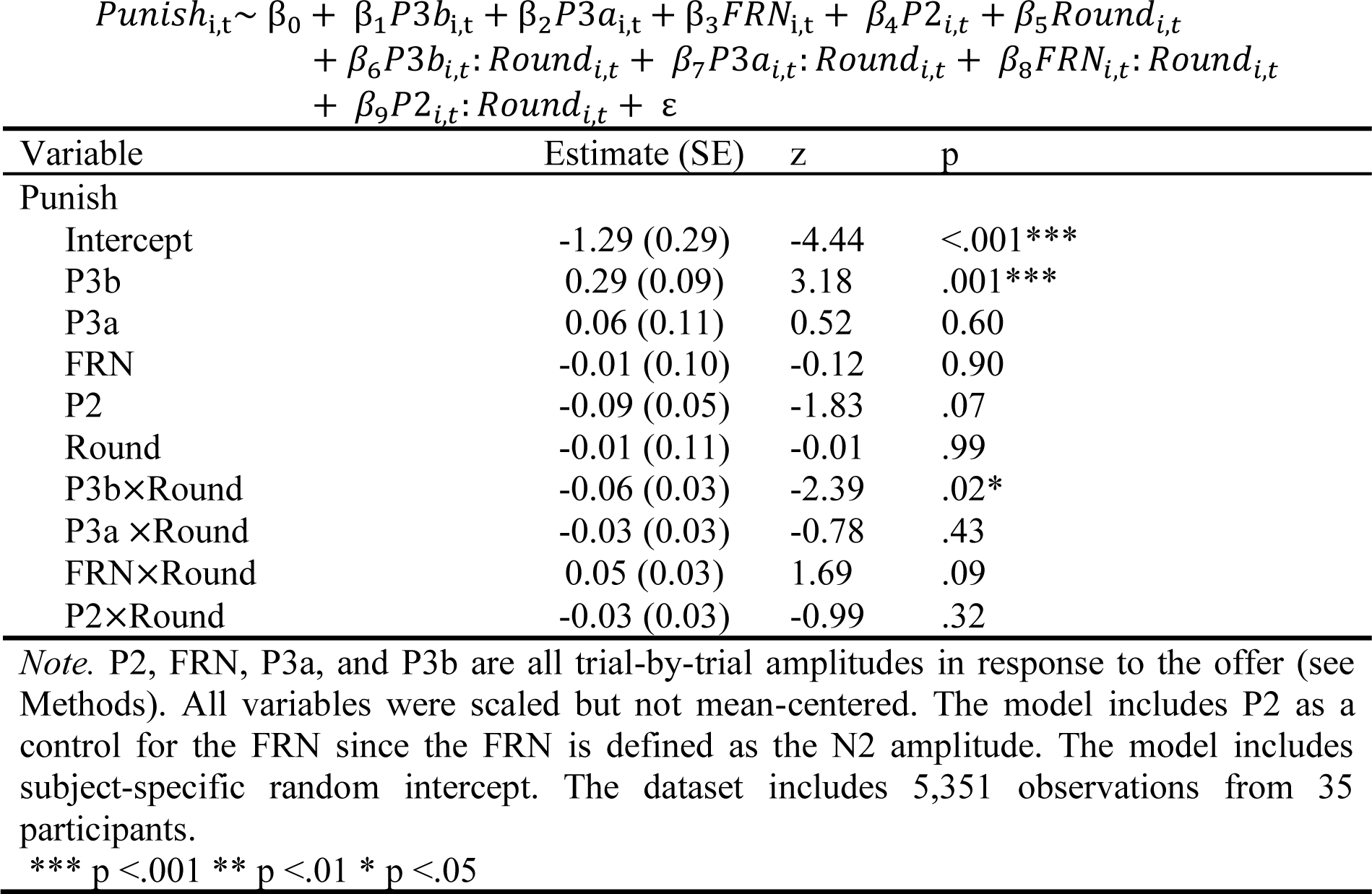
Only P3b predicts choices over rounds.

### Reward and emotion PEs indexed by separate and dissociable neural signals

We next investigated if each PE—reward, valence, and arousal PEs—is represented by distinct and separable neural architecture. We first standardized all PEs at the group level and used each PE type as independent variables in separate linear mixed effects regressions predicting our *a priori* ERPs of interest: the FRN, P3a, and P3b (see Methods). The FRN is known to encode both signed and unsigned PEs^22,29^, and RPE effects on the P3 component are also somewhat ambiguous on what they index^30^, potentially reflecting the magnitude of the PE rather than the valence^23^. Given the lack of clarity around what exactly these ERPs index, we first tested whether absolute or signed PEs predicted the ERPs of interests, and found more evidence for absolute, compared to signed, PEs (see Supplement). Using the absolute value of each PE as the predictor, we found that trial-by-trial FRN amplitudes were uniquely predicted by reward PEs (β = −0.23 ± 0.07, *t* = −3.17, *P* = 0.003) but not valence (β = 0.06 ± 0.07, *t* = 0.91, *P* = 0.36) or arousal PEs (β = 0.04 ± 0.08, *t* = 0.44, *P* = 0.67; Fig. 3). In contrast, trial-by-trial P3b amplitudes were only predicted by valence PEs (β = 0.06 ± 0.02, *t* = 3.98, *P* < 0.001), but not arousal (β = −0.001 ± 0.01, *t* = −0.71, *P* = 0.48) or reward PEs (β = 0.03 ± 0.02, *t* = 1.37, *P* = 0.18; *Fig. 3*), and a beta-coefficient test showed that the valence PE effect was marginally greater than the reward PE (*Z* = 1.44, *P* = 0.07). Finally, trial-by-trial P3a amplitudes were only weakly linked to arousal PEs (β = 0.04 ± 0.02, *t* = 1.90, *P* = 0.07), and not associated with either valence (β = 0.01 ± 0.02, *t* = 0.18, *P* = 0.86) or reward PEs (β = 0.04 ± 0.02, *t* = 1.69, *P* = 0.10; Fig. 3; see supplement for an exploratory model showing that the P3a is best tracked by offer extremity). To ensure that we were capturing all possible electrophysiological signatures of reward, valence, and arousal PEs (which we might have missed with a pure *a priori* ERP approach) we additionally employed a data-driven method that does not rely on predefined signals^31^. Results from this data driven approach showed converging evidence for separate spatiotemporal clusters in response to offers for reward and valence PEs (see Supplement). Taken together, these results are the first to illustrate that emotion and reward learning signals are separately encoded in the brain.

**Figure 3.**
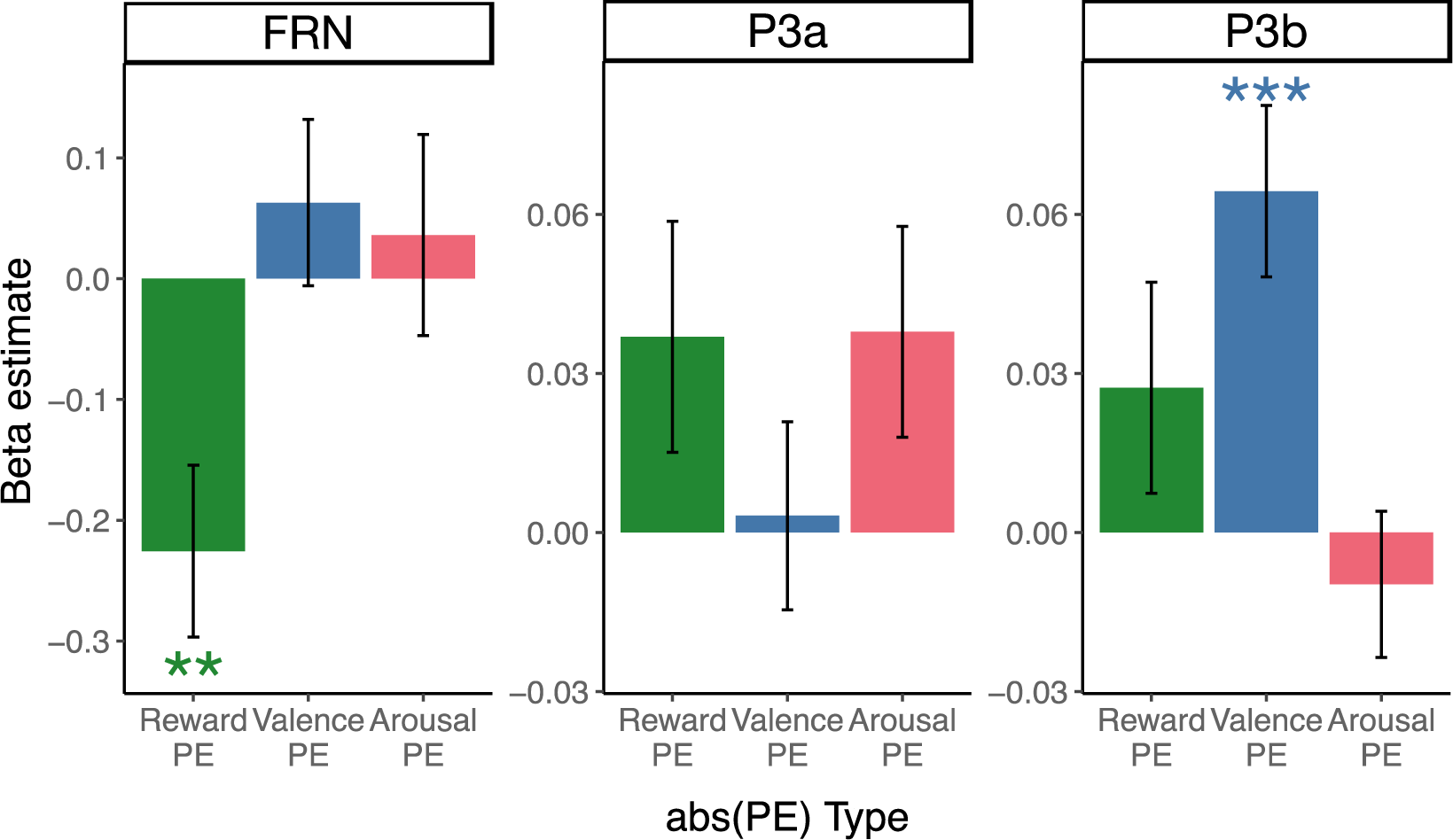
Separate ERPs index reward and affective PEs. The data on each graph reflect the beta coefficient from LMMs modeling the marginal contributions of the absolute value of each PE type on separate ERPs: the FRN, P3a, and P3b. Error bars reflect ±1 S.E. ***P<.001, **P<.01,

To assess the neural learning effects on choice, especially given our behavioral findings showing that the relationship between PEs and choice varies with round number, we next allowed each ERP to interact with round. Results reveal a significant simple effect of P3b on choice (Table 2), as well as an interactive effect between P3b and round on choice (Table 2; Fig. 2B). This mirrors our behavioral finding, revealing that the P3b—which is uniquely associated with valence PEs—is the sole neural signal driving choice, and the strength of this neural signal diminishes over time as more information is gleaned about a partner’s behavior. Although not statistically significant, the interaction between FRN and round also reflects the behavioral pattern, where the relationship between FRN and choice strengthens with time (Table 2; Fig. 2B). Collectively, these findings illustrate that both FRN and P3b uniquely track reward and emotion learning signals, but that only emotion PEs, indexed by the P3b, are relevant for choice.

### Reward and affective PEs are resolved through different mechanisms

Although most PEs are no longer predictive by the final round—indicating rapid learning—it remains unclear which component of the PE drives the error signal. On one hand, participants might adjust their expectations to make upcoming events less surprising. This dovetails with reinforcement learning accounts which predominantly emphasize adjusting PEs by altering expectations (i.e., increasing Q-value of an action to anticipate a greater reward next time). On the other hand, participants could alter experiences (perhaps by employing emotion regulation tactics) to lessen an event’s impact. While research on affective forecasting suggests that accurately predicting future emotional events is challenging^32,33^, one could use emotion regulation strategies to modify responses to events like unfair offers^34^. These two accounts present divergent theories about how affective and reward PEs might drive learning.

To explore the theory that modifying expectation can reduce both affective and reward PEs, we examined how reward, valence, and arousal expectations changed throughout the task. As expected, participants show the largest update in both reward and emotion expectations between the first and second rounds (Table 3, Fig. 4). We then probed how expectations change between rounds two through five to understand if participants continue to adapt after an initial surprise. Reward expectations for this period reveal that participants continue to refine beliefs about their partners, baring evidence of continued learning through reward PEs. In contrast, expectations about valence and arousal remain largely consistent across all partner types from rounds two through five, suggesting participants do not continue to adjust their affective expectations after the initial round. In other words, only reward—and not affective—PEs are resolved by altering expectations. Next, we examined whether experiences change across the task. We found that participant’s subjective reports of their valence and arousal experiences changed significantly over rounds. Specifically, negative reactions to unfair offers wane over time, and the intense positive feelings of receiving a fair offer also diminish as you learn more about the fair partner (Table 3, Fig. 4). Given that monetary offers are fixed by task design, offers do not significantly vary across rounds. Thus, these results paint a clear distinction in how affective and reward PEs are leveraged for learning: after an initial large update of expectations, reward PEs are resolved by adjusting reward expectations (dovetailing with the RL literature), whereas affective PEs are managed by aligning emotional experiences with prior predictions.

**Figure 4.**
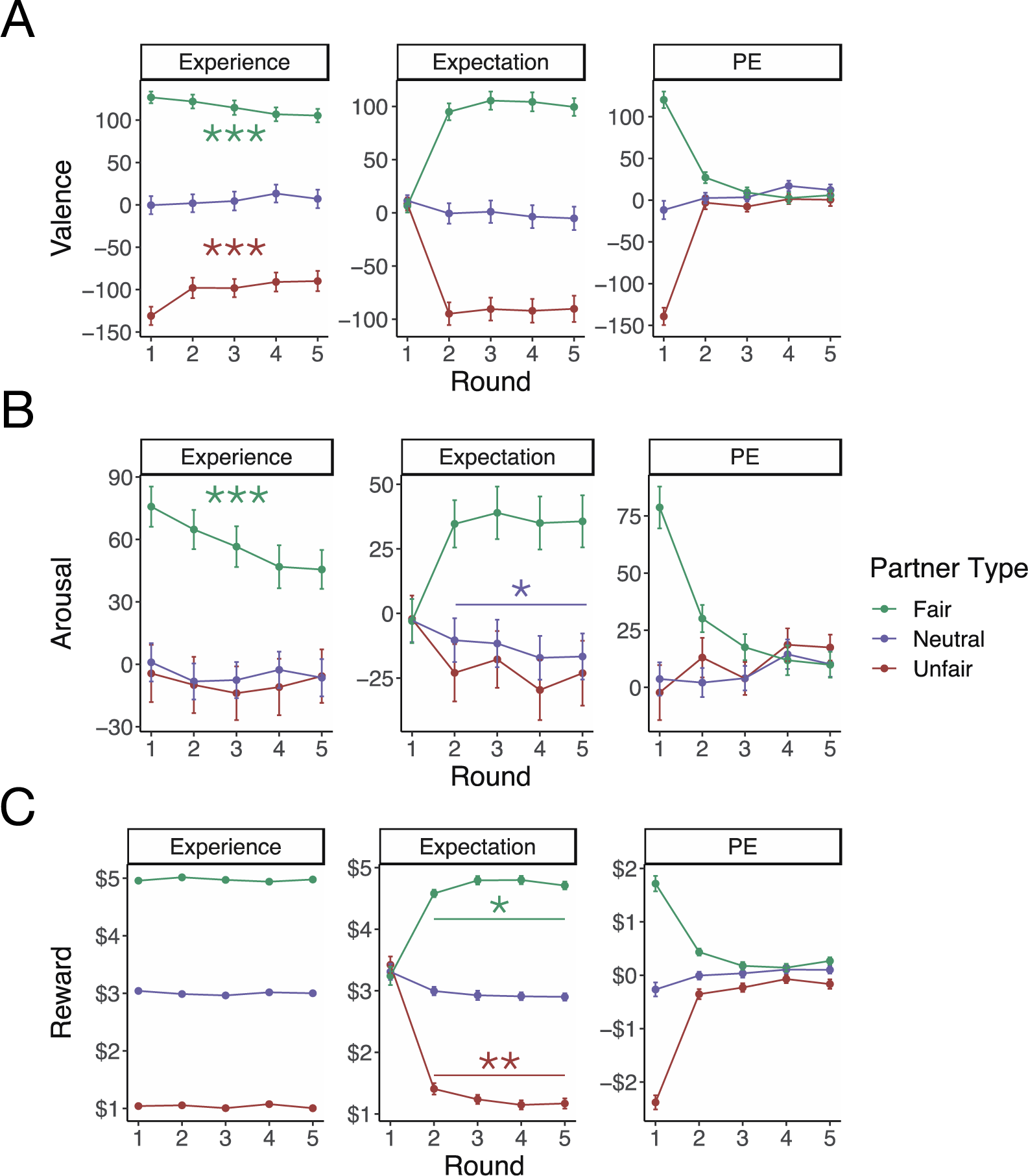
Learning in the Ultimatum Game. **A) Valence measurements.** Valence PEs are calculated as the difference between valence experience and expectations for each round. Valence ratings are between -250 (unpleasant) and 250 (pleasant). **B) Arousal measurements**. Arousal PEs are calculated as the difference between arousal experience and expectations for each round. Arousal ratings are between -250 (low intensity) and 250 (high intensity). **C) Reward measurements**. Reward prediction errors (PEs) are calculated as the difference between reward offer and reward expectations for each round. Reward measurements are between $0 and $10.All data are averaged within and then across participants for each round and partner type. All error bars reflect ± 1 S.E. Statistical tests are shown in Table 2. ***P<.001, **P<.01, *P<.05.

**Table 3.** Different mechanisms underlying resolution of reward and affective Pes.

| | Round 1 – 2 Update<br>(Expectations)<br>*t*-score (*P*-value) | Rounds 2-5 Update<br>(Expectations)<br>$\beta \pm SE$ (*P*-value) | Round 1 – 5 Update<br>(Experience)<br>$\beta \pm SE$ (*P*-value) |
| --- | --- | --- | --- |
| **Fair partner** |  |  |  |
| Reward | 9.53 (<.001***) | 0.02±0.01 (.02*) | - |
| Valence | 9.23 (<.001***) | 0.01±0.01 (.44) | -0.04±0.01 (<.001***) |
| Arousal | 5.00 (<.001***) | -0.001±0.02 (.94) | -0.08±0.02 (<.001***) |
| **Neutral partner** |  |  |  |
| Reward | -3.00 (.005**) | -0.02±0.01 (.09) | - |
| Valence | -1.44 (.16) | -0.02±0.01 (.16) | 0.02±0.01 (.05) |
| Arousal | -1.45 (.15) | -0.03±0.01 (.02*) | -0.01±0.01 (.35) |
| **Unfair partner** |  |  |  |
| Reward | -14.19 (<.001***) | -0.05±0.02 (.006**) | - |
| Valence | -10.36 (<.001***) | 0.01±0.02 (.51) | 0.07±0.02 (<.001***) |
| Arousal | -2.28 (0.03*) | -0.01±0.02 (.55) | -0.003±0.01 (.82) |

*Note.* Rounds 1-2 update (expectations) shows the result of a paired *t*-test comparing expectation values from round 2 to round 1. Rounds 2-5 update (expectations) shows the result of a LMM comparing how expectation values change between rounds 2-5. Rounds 1-5 update (experience) shows the result of LMMs comparing how emotional experiences change between rounds 1-5. Reward experiences were defined by task parameters and are stable across rounds (see Methods). \*\*\* $P < .001$ , \*\* $P < .01$ , \* $P < .05$ .

## Discussion

While emotions clearly influence learning and decision-making^35^, most reinforcement learning models do not incorporate emotions as an error signal driving social learning. Instead, these models predominantly emphasize reward PEs as the central driver of behavior, with some exceptions. In this study, we leverage EEG to determine if emotions serve as a key learning signal in a repeated economic game, and whether these emotion signals can be differentiated from reward PEs. Our behavioral results show that affective PEs have an independent and stronger contribution to choice, especially when there is significant uncertainty about a partner’s actions. As this uncertainty decreases with experience, reward PEs begin to play a more dominant role in guiding choice. At the neural level, distinct ERPs were found to index each PE: the FRN corresponds most closely with reward PEs, the P3b is associated with valence PEs, and the P3a is indexed by arousal about offer extremity. Collectively, these findings suggest that reward and emotion learning signals have dissociable neural locations with distinct temporal trajectories. At the trial level, that the temporal trajectory of the FRN comes online first and is then followed by the P3b, suggests that monetary offers are likely initially evaluated based on how surprising the reward is (e.g., “How does this monetary offer differ from my reward expectations?”) and only later are violations of emotion expectations incorporated (e.g., “I feel better/worse than anticipated”). An examination of social learning over time, however, shows that affective PEs are most powerful during early trials and attenuate as uncertainty is reduced. In contrast, reward PEs follow the opposite pattern, growing in predictive power over time.

In contrast to reward-centric accounts that dominate the learning literature^38^, our findings underscore that violations of affect expectations play a particularly privileged role in social learning, a role that cannot be solely subsumed by rewards. Although rewards may be sufficient for building successful artificial agents, there is growing concern about whether external reward functions are enough to explain the breadth and flexibility of human decision-making, and it has been suggested that emotional signals might bridge this gap^39–41^. While prior affective research has primarily explored long-term affective forecasting errors^33^ or contexts absent of trial-by-trial learning^19^, here we provide a direct assessment of affective learning signals at the neural level. Specifically, we find that the P3b component, which tracks valence PEs, stands as the sole neural predictor of social choice. This aligns with prior work showing that the P3b integrates information from multiple learning mechanisms relevant for decision policy changes^31,42^. While far less influential, reward PEs still contribute to learning^22,29,43,44^, and our findings show that the FRN uniquely indexes monetary reward PEs—even when controlling for affective signals. In summary, a comprehensive explanation of human decision-making necessitates consideration of both rewards and emotions.

While research indicates that emotional stimuli, such as faces or words, are sometimes processed relatively early^45–47^, our results indicate that evaluations of emotionally charged social exchanges are initially evaluated based on reward. While at first blush this might appear as a discrepancy, emotional experiences unfold over time, which include initial attention to the event and subsequent evaluation^48^. That we probe participants’ self-reports of their affective experience following the offer, may align more consistently with the evaluation of an affective experience rather than initial attention. This account aligns with other EEG work showing that late neural components, such as the Late Positive Potential (LPP), are associated with emotional stimuli^49^. Interestingly, the LPP shares similar morphology with the P3b, is sensitive to a stimulus’s emotional saliency^50^, can be influenced by cognitive reappraisal or attentional shifts^51^, and is also known to modulate attention during late-stage processing^52,53^. Taken together, this suggests that emotional processing that unfolds on later temporal trajectories can still be influential for higher cognition.

Our data also suggests that affective experiences habituate with repeated experiences, giving rise to smaller prediction errors over time. The consequence of blunted affective responses through experience (i.e., reduced prediction errors) means that the emotional experiences of an initial interaction with a social partner is amplified compared to subsequent encounters. This pattern is consistent with normative prescriptions for learning in uncertain and dynamic environments^54–56^, highlighting the possibility that the attenuation of affective experience might serve an important role in optimizing learning under uncertainty. Future work could explore this further by extending our paradigm to manipulate a broader array of factors that influence normative learning dynamics, such as outcome stochasticity^55,57^, volatility^54,57^, and temporal structure^31,58^.

By taking the simple, albeit novel, step of incorporating emotion as an error signal into a framework, we reveal the pivotal role of affective PEs in driving social learning. With the precise temporal neural time course of EEG, we provide evidence for early processing of reward PEs and the later processing of affective PEs when learning about other people. Although violations of emotion expectations are integrated relatively late in the decision process, they play the strongest role in predicting social choice—providing evidence of a neurobiologically plausible distinct emotion error signal.

## Materials and methods

### Participants

Participants (N = 41, 25 female, mean age = 20.8 ± 4.4) received either monetary compensation ($15 per hour) or course credits and provided informed consent in a manner approved by Brown University’s Institutional Review Board under protocol 1607001555. A power analysis of the unique effect of valence prediction error on choice in our prior work^19^ revealed that 18 participants would be sufficient to detect this effect with an alpha of 0.05 and power (beta) of 0.80. Accordingly, we aimed to exceed this and collected a sample of 40 participants which matches sample sizes of recent EEG studies focusing on the FRN and P300^31,42^.

### Task and procedure

Participants played an adapted repeated Ultimatum Game^59,60^ that included subjective emotion and reward ratings^19,28^. Participants were told they were playing with past participants who gave offers across five rounds, conditional on participant’s choices to accept or reject each offer—similar to strategy methods used in economics^61^. Unknown to participants, their partner’s offers were generated from one of three normal distributions representing three types of proposers: 1) “unfair” proposers gave offers according to a normal distribution with a mean of $1 and SD of 0.50; 2) “neutral” proposers gave $3 on average with a SD of 0.50; and 3) “fair” proposers who gave $5 on average with a SD of 0.50. Participants played with 36 unique partners, 12 of each type and all five offers from each partner were randomly drawn from their respective normal distribution. We used faces from the 74 image MR2 database^62^ to represent partners and 36 faces were pseudo-randomly pulled from this database per participant to achieve a balanced distribution of images of men and women of European, African, and East Asian ancestry. Subjective affective predictions and experiences were reported using a 500-500 pixel two-dimensional affect grid where the horizontal axis was valence (unpleasant/pleasant feelings) and the vertical axis was arousal (low/high intensity feelings). Both dimensions range from -250 to +250. To familiarize participants with this affect grid, participants completed an emotion classification task prior to the repeated UG. Participants made affect ratings of 20 canonical emotion words (for example, angry, sad, and surprised) on the grid, twice for each word, in a randomized order. Training participants to interpret this subjective affect grid has shown strong convergent validity with other approaches for emotion ratings^63^.

We calculated affective prediction errors (PEs) on a trial-by-trial basis by measuring the discrepancy between participants’ actual affective experiences and their affect expectations. Emotion PEs can be defined on both valence and arousal dimensions. A valence PE was computed by subtracting the predicted level of (un)pleasantness of an offer from the actual experienced (un)pleasantness, while an arousal PE was the difference between the expected arousal and actual experienced arousal. For instance, if a participant felt unpleasant about receiving an offer (e.g., rating it -200) but had anticipated feeling slightly pleasant (e.g., rating it +40), the valence PE would be -240 (-200 minus +40). Similarly, reward PEs were calculated by subtracting the predicted monetary reward from the actual offer given to the participant for each trial.

The was UG comprised of nine blocks of four partners each (20 trials per block), with self-paced rests between blocks for a total of 180 trials. Participants additionally completed pre-UG and post-UG likability ratings for all 36 partners on a visual analog scale (0 – 10, in increments of .01). The experiment was delivered in Matlab (The MathWorks, Inc.) using the Psychtoolbox-3 package and included stimulus presentation, event, and response logging. A standard computer mouse and keyboard were used for response registration.

During the UG, participants were first shown a picture of their partner for 1000ms, followed by a fixation cross (500ms, same timing for all fixations). Participants were then given cues for reward predictions ($?) or affect predictions (E?; 1000ms each); these cues indicate that participants will be making the required response on the next screen. Reward predictions (how much participants expect the partner to offer) are reported on a visual analog scale ($0 - $10) and participants have unlimited time to respond. Affect predictions (how participants expect to feel after the offer) are reported on the valence-arousal grid and participants are required to answer within 5s. The order of predictions was counterbalanced and separated by fixations. Following predictions, the offer was given (2000ms) in dollar and cent format (e.g., $2.37), and followed by another fixation. Participants were then given an affective experience rating cue (E; 1000ms), which indicated that they should rate how they felt about the offer using the affect-grid (required within 5s). A fixation followed and then participants were given a choice cue (C; 1000ms) indicating they would need to make their choice. The choices to accept or reject were presented on the screen (e.g., [A] [R]), and the order of these options was counterbalanced. When participants were matched with a new partner, they were presented with a waiting screen for 1-4s before starting the next trial.

Prior to experiment, participants filled in the following two personality questionnaires: the 20-item Toronto Alexithymia Scale^64^ and the Temporal Experience of Pleasure Scale^65^. These measures were registered as potential control variables and for other purposes not addressed here. Participants were then seated in a shielded EEG cabin. Prior to completing the emotion classification and UG task, participants performed practice trials.

### Psychophysiological recording and processing

EEG was recorded using BrainVision recorder software (Brain Products, München, Germany) at a sampling rate of 500 Hz from 64 Ag/AgCl electrodes mounted in an electrode cap (ECI Inc.). Data was collected using Cz as a reference channel and re-referenced to average reference offline. Electrodes below the eyes (IO1, IO2) and at the outer canthi (LO1, LO2) recorded vertical and horizonal ocular activity. At the end of the experiment we recorded prototypical eye movements (20 trials of each: up, down, left, and right) for offline ocular artifact correction. We kept electrode impedance below 10 kΩ.

EEG data were processed using Matlab (The MathWorks Inc.) using the EEGlab toolbox^66^ as previously described^42^ and included the following steps: (1) re-referencing to average reference and retrieving the Cz channel, (2) removal of blink and eye movement artifacts using BESA^67^, (3) bandpass filtering of .1 – 40 Hz, (3), (4) epoching the ongoing EEG from -200 to 800ms relative to offer onset, (5) removal of segments containing artifacts, based on values exceeding ±150 µV and gradients larger than 50 µV between two adjacent sampling points. Baselines were corrected to the 200ms pre-stimulus interval (offer onset) using the regression method in subsequent analyses^68^.

To define the time windows for single-trial analyses of FRN, P3a and P3b amplitudes, we first determined the grant average peak latencies of FCz, FCz, and Pz, respectively. Accordingly, the FRN was quantified on single trials as the average voltage within an interval from 315 to 415ms after offer onset across all electrodes within a fronto-central region of interest including F3, Fz, F4, FC3,FCz, FC4, C3, Cz, C4^23^. To control for P2 effects on the FRN, the P2 amplitude was also extracted within each trial as the average voltage between 199-299ms across fronto-central electrodes F1, Fz, F2, FC1, FCz, FC2, C1, Cz, C2, and included as a regressor in the analyses. P3a amplitude was quantified on single trials as the average voltage within a 363-463ms interval post-offer across fronto-central electrodes F1, Fz, F2, FC1, FCz, FC2, C1, Cz, C2^42^. P3b amplitude was quantified on single trials as the average voltage within a 530-630ms interval post-offer within a parietally-focused region of interest including CP1, CPz, CP2, P1, Pz, P2, PO3, POz, PO4^42^.

### Analyses

Reward PEs (RPE) was determined as the offer minus the reward prediction given by participants Valence and Arousal PEs were determined similarly: the affective experience participants reported upon receiving the offer minus participant’s affective prediction for how they would feel after the offer. Prior to analyses, reward, valence, and arousal PEs were standardized but not mean centered, as zero represents a meaningful value on these scales (predicted and actual experiences are the same). Inspection of the behavioral data identified four trials in which impossible affect ratings were given (valence or arousal ratings outside of the 500 by 500-pixel grid) and these data were excluded from relevant analyses.

## Notes

### Competing Interest Statement

The authors have declared no competing interest.

