## Supplement for "Dissociable neural signals for reward and emotion prediction errors"

**Contents:**

1. Supplementary Methods
2. Supplementary Results
3. Supplementary References

### 1 Supplementary Methods

#### 1.1 Task Instructions

Participants completed an Emotion Classification (EC) task before playing the Ultimatum Game (UG). In the EC, participants rated 20 feeling words two times each on an affect grid used in past research[1,2]. Words were presented in a randomized order. Instructions for both are presented below:

**<u>Instructions for the Emotion Classification Task</u>**

*The purpose of this task is to study how people classify feelings. In this experiment, you will use diagrams like the one to the right to describe feelings. It is in the form of a square – a kind of map for feelings. The center of the square represents a neutral, average, everyday feeling. It is neither positive nor negative.*

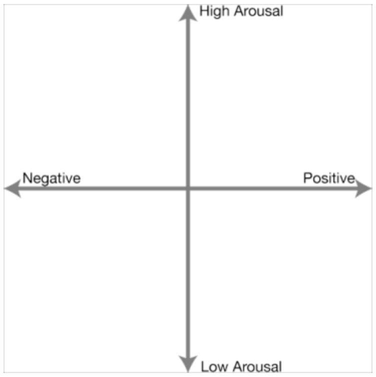


*The right half of the grid represents pleasant feelings. The farther to the right the more pleasant.  The left half represents unpleasant feelings. The farther to the left, the more unpleasant.*

*The vertical dimension of the map represents degree of arousal. Arousal has to do with how wide awake, alert, or activated a person feels – independent of whether the feeling is positive or negative. The top half is for feelings that are above average in arousal, and the lower half for feelings below average. The bottom represents sleep, and the higher you go, the more awake a person feels. At the top is maximum arousal. If you imagine a state we might call frantic excitement (remembering that it could be either positive or negative), then this feeling would define the top of the grid.*

*In this part of the study, you will be briefly shown a feeling word in the center of the screen. Try to think about what the feeling is like. Some people think about situations, or draw on memories of situations, that have made them feel that in the past. After seeing the word, the next screen will show the diagram to the right and ask you to rate the feeling.*

*You can click on the diagram to rate the feeling only once so make sure of your response before pressing the mouse button.*

**Instructions for the Ultimatum Game**

*The purpose of this task is to study how people make decisions. You will be making real decisions that affect the monetary outcomes of YOURSELF and OTHERS. You will be playing 5 rounds in a row with a partner before being matched with different partners.*

*Your partner has been allotted $10. Your partner has already decided how much of their $10 to offer you. Your partner can offer you any amount of their $10, ranging from nothing ($0) all the way to everything ($10) You have two choices in response to your partner's offer:*
*You can accept the offer and keep both you and your partner's money the same*
*You can reject the offer and decrease both you and your partner's money to 0*

*Ultimately, you will decide how much money you and your partner actually receive.*

*Each round will be broken up into different stages. It is very important that you learn the cues associated with each stage, which we will go over now.*

*Partner Face: At the beginning of each trial, you will see a picture of your partner. It is important that you notice and attend to them*

*Reward Prediction (?): This cue is a dollar sign followed by a question mark. This cue indicates that you will be asked to think about and rate how much you think your partner will offer you, ranging from $0 to $10 using your mouse*

*Emotion Prediction (?): This cue is a capital letter "E" followed by a question mark. This cue indicates that you will be asked to think about and rate you think you will feel when you receive the offer from your partner. You will use the emotion grid to make your rating with your mouse.*

*Sometimes the emotion prediction will be first. Other times the reward prediction will be first.*

*Offer ($X.XX): This is not a cue, but displays your partner's offer for that round in the following format. It is important that you notice and attend to the offer amount, since it will only appear for a brief time at the center of the screen.*

*Emotion Experience (E): This cue is the capitalized letter "E". This cue indicates that you will be rating how you feel about the offer using the emotion grid with your mouse.*

*Choice (C): This cue is the capitalized letter "C". This cue indicates that you will be making your choice about your partner's offer. You can either accept the offer or reject the offer. The options to accept or reject will appear in the center of the screen with brackets such that [A] will be accept and [R] will always be reject.*
*IMPORTANT: the order of these options can change ([A] [R] or [R] [A]) so it is very important that you notice which side the option is on. You will use the buttons D to select the left option and F to select the right option.*

*You are playing with past participants who came into our lab and completed this task. Just like you will be asked, they were asked to make economic choices, which were recorded and put into a database which we will use today. They were photographed and asked to decide how much of their $10 to offer future participants in this game. Critically, they did this for 5 rounds based on every possible choice you could make. Their responses are what you are seeing today. Therefore, you will be interacting with past volunteers, and your decisions will impact not only your monetary payment but their monetary payment as well.*

*To determine the final payouts of both players, the program will randomly select 1 partner at the end of the experiment. All 5 trials with that partner will be realized. That is, both you and your partner will receive an additional BONUS for those decisions. This bonus is in addition to the $45 you will make just for participating in the study. We will pay you today and your partner through Venmo.*

### 2 Supplementary Results

#### 2.1 Behavioral Results

*Learning about social partners changes how an offer is evaluated*. What impact does learning about social partners have on participants' choices? Using generalized mixed effects linear regression, we found that across rounds participants rejected unfair offers (57%) more often than neutral offers (29%; $\beta = 2.09 \pm 0.22, z = 9.46, P < .001$), and in turn rejected neutral offers more often than fair offers (8%; $\beta = 2.49 \pm 0.38, z = 6.60, P < .001$; Fig. S1A). When we examined choices over rounds, we found that the probability of rejecting offers from unfair $(\beta = -0.16 \pm 0.04, z = -4.31, P < .001)$, and neutral $(\beta = -0.08 \pm 0.04, z = -1.91, P = 0.06)$ partners decreased over rounds. This suggests that participants might be changing their decision policy as rejections near the end of the game have no strategic power to influence future offers. Interestingly, rejection of offers from fair partners actually increased over rounds ($\beta = 0.22 \pm 0.06, z = 3.69, P < .001$; Fig. S1B). One possible explanation for these change in policy is that participants' evaluation of offers is conditional on partner type. Figure S1C shows the distribution of unfair, neutral, and fair partners' offers, which have partial overlap between certain partner types (unfair and neutral, neutral and fair). Generalized linear mixed-effects models showed main effects for partner type, indicating that the same offer (e.g., $2.15) is rejected more if it came from a neutral partner than if it came from an unfair partner $(\beta = 1.53 \pm 0.60, z = 2.54, P = .01)$. A similar effect existed between neutral and fair partners, such that the same offer (e.g., $3.50) is rejected more from fair partners compared to neutral $(\beta = 3.94 \pm 1.37, z = 2.89, P = .004)$. In other words, participants evaluated current offers relative to their general expectations for each partner (unfair: $1, neutral: $3, fair: $5), and these expectations changed how they evaluated offers and made decisions, consisting with theories about the efficient coding of subjective value[3].

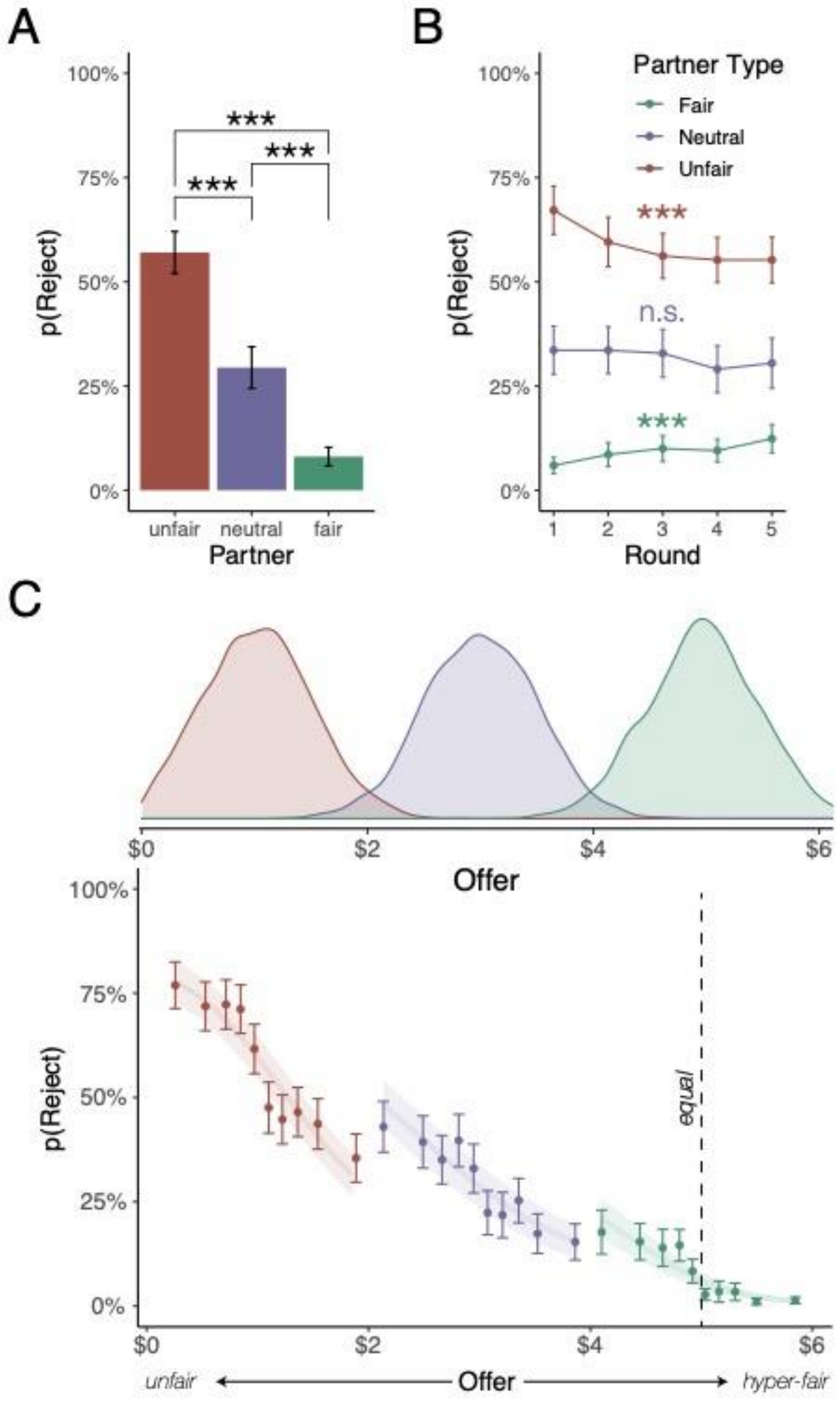
A
B
C
p(Reject)
100%
75%
50%
25%
0%
***
unfair
neutral
fair
Partner
Partner Type
Fair
Neutral
Unfair
n.s.
1
2
3
4
5
Round
$0
$2
$4
$6
Offer
equal
unfair
hyper-fair

***Figure S1. Choice behavior in the Ultimatum Game.*** ***A)*** *Overall choices per partner type. Bars reflect average proportion of rejections per partner type.* ***B)*** *Choices over round per partner type. Dots reflect average proportion of rejections per round per partner type. Analyses reflect regression effects.* ***C)*** *Offer evaluation by partner type. Histograms reflect empirical distribution of offers for partner type across participants. The probability of rejecting the offer is plotted for all three partner types and analyses represent regression effects. All error bars and shaded areas reflect ±1 S.E. ***P<.001, **P<.01, *P<.05.*

*Reward and emotion are updated separately during learning*. While results from the manuscript show that valence and reward PEs independently contribute to decisions to punish, it is possible that these PEs contain redundant information for learning. To test whether each PE is updated separately, we ran separate LMMs modeling updates for valence, reward, and arousal expectations between rounds, defined as the difference between expectations on *t and t-1*. For example, if reward PE contains all the necessary information to predict the update for valence expectations, then reward PEs should exhibit the strongest contribution during valence updating. We did not find this to be the case. Instead, updating for each PE type is strongly driven by the information stored in its own PE, such that reward PEs predict reward updating ($\beta_{reward\ PE} = 0.32 \pm 0.01, t = 41.30, P < 0.001$), valence PEs predict valence updating ($\beta_{valence\ PE} = 0.57 \pm 0.03, t = 21.17, P < 0.001$), and arousal PEs predicts arousal updating ($\beta_{arousal\ PE} = 0.48 \pm 0.03, t = 17.01, P < 0.001$; *Fig. S2*). These results suggest that each PE provides some unique information that drives updating expectations during trial-by-trial learning.

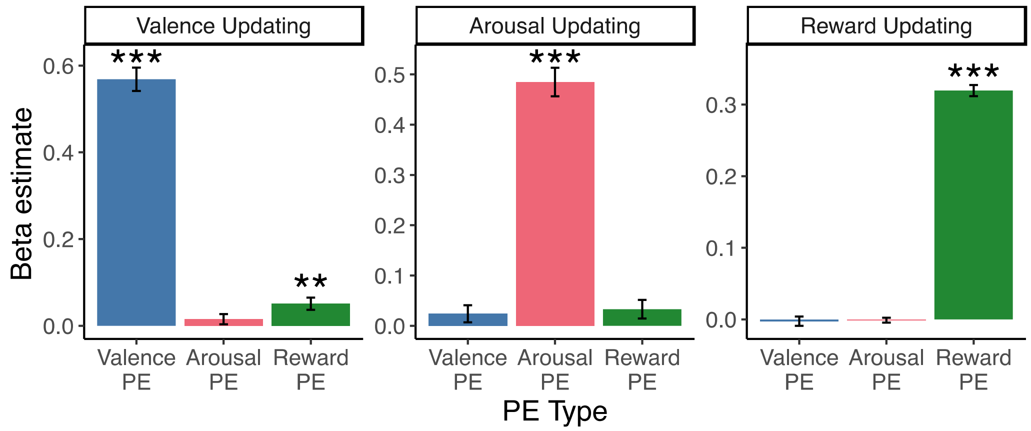


***Figure S2. Separate updating of valence, arousal, and reward PEs.*** *The data on each graph reflect the beta coefficient from LMMs modeling the expectation of the future round (i.e., updating) as a linear combination of all three PE values on the current round. Higher positive values reflect stronger contribution to updating. Error bars reflect ±1 S.E. ***P<.001, **P<.01, *P<.05.*

#### 2.2 Neural Results

*Waveforms of the FRN, P3a, and P3b in the repeated UG.* The exact timing of a waveform can differ across experiments, and to determine when the FRN, P3a, and P3b occurred we calculated the average peak latencies at the FCz, FCz, and Pz, respectively. The FRN was quantified as the average voltage within an interval from 315 to 415ms after offer onset. To visualize this waveform, we averaged across all electrodes within a fronto-central region of interest including F3, Fz, F4, FC3, FCz, FC4, C3, Cz, C4[4] for each partner type (Fig. S3). The P3a was quantified as the average voltage within a 363-463ms interval post-offer. To visualize this waveform, we averaged across fronto-central electrodes F1, Fz, F2, FC1, FCz, FC2, C1, Cz, C2[5] for each partner type (Fig. S4). Lastly, the P3b was quantified as the average voltage within a 530-630ms interval post-offer. To visualize this waveform, we averaged across a parietally-focused region CP1, CPz, CP2, P1, Pz, P2, PO3, POz, PO4[5] for each partner type (Fig. S5).

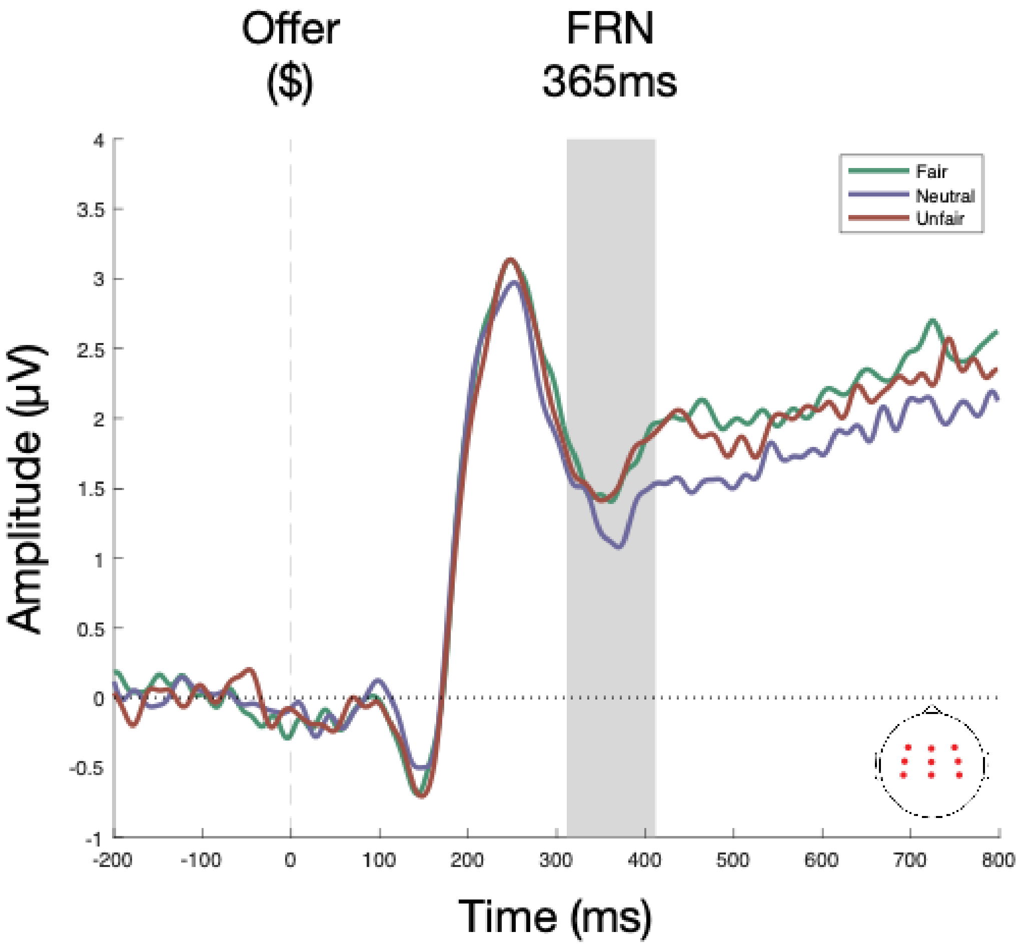


***Figure S3. FRN waveform.*** *Amplitudes across the depicted electrode topography were first averaged within subject, then across subjects for each partner type. The shaded bar shows the interval for the FRN defined by mean subject-wise peak latencies at the FCz.*

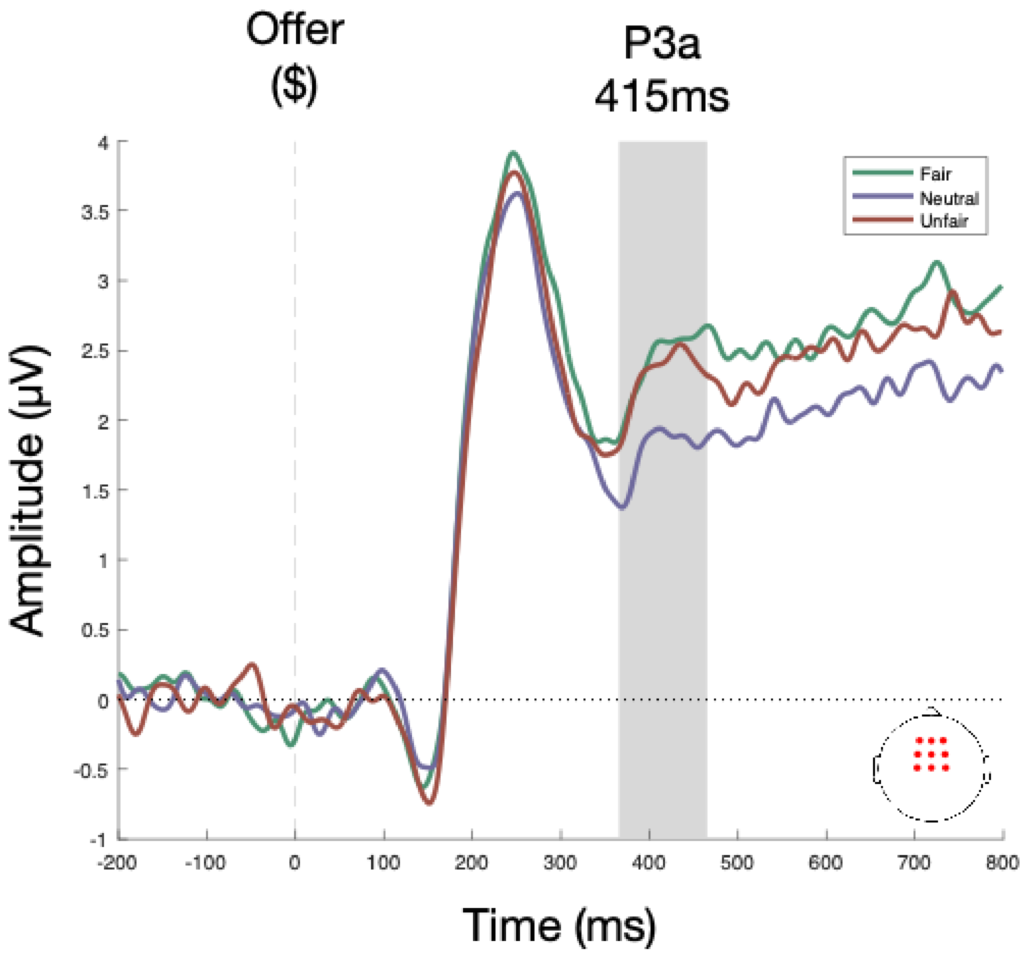


***Figure S4. P3a waveform.*** *Amplitudes across the depicted electrode topography were first averaged within subject, then across subjects for each partner type. The shaded bar shows the interval for the P3a defined by mean subject-wise peak latencies at the FCz.*

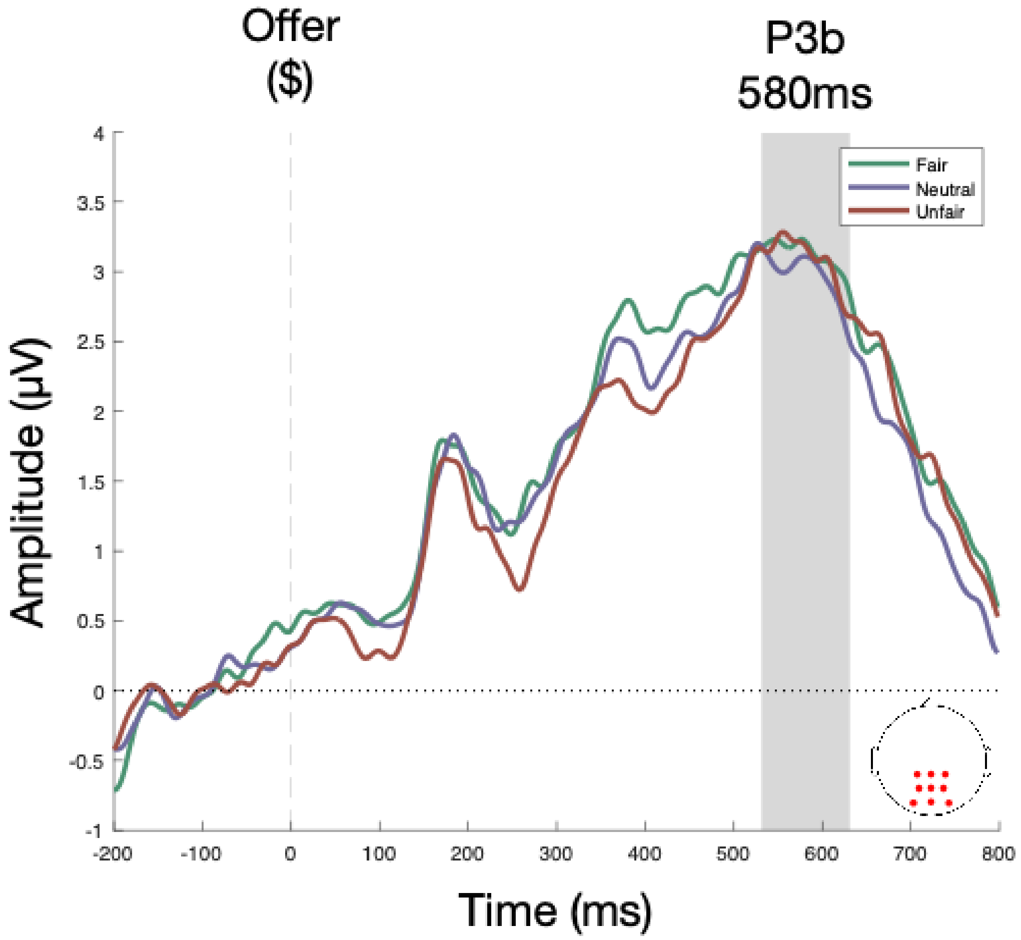


***Figure S5. P3b waveform.*** *Amplitudes across the depicted electrode topography were first averaged within subject, then across subjects for each partner type. The shaded bar shows the interval for the P3b defined by mean subject-wise peak latencies at the Pz.*

*Signed PEs largely do not relate to ERPs of interest.* There is a significant ongoing debate whether EEG can be used to detect both signed and unsigned reward PEs[6]. Accordingly, we tested whether the Feedback-Related Negativity (FRN) exhibited sensitivity to signed reward, valence, and/or arousal PEs. Results showed that trial-by-trial FRN amplitudes were not predicted by signed reward PEs ($\beta = 0.05 \pm 0.06, t = 0.74, P = 0.47$), valence PEs ($\beta = -0.11 \pm 0.06, t = -1.87, P = 0.07$), nor arousal PEs ($\beta = -0.003 \pm 0.04, t = -0.07, P =$

0.95). While the P300 effects are typically associated with unsigned PEs[4], we extended our analysis of signed PEs to test whether the P3a and P3b components are sensitive to our signed predictors. Trial-by-trial P3a amplitudes were strongly predicted by signed arousal PEs ($\beta = 0.03 \pm 0.01, t = 3.21, P = 0.001$) but not valence PEs ($\beta = -0.007 \pm 0.02, t = -0.44, P = 0.66$) or reward PEs ($\beta = -0.01 \pm 0.01, t = -0.81, P = 0.42$). This finding is intriguing because arousal PEs, reflecting variations in arousal compared to expectations, serve as saliency signals and are inherently non-valenced. Lastly, trial-by-trial P3b amplitudes were predicted by reward PEs ($\beta = -0.03 \pm 0.01, t = -2.53, P = 0.01$) but not by valence PEs ($\beta = -0.009 \pm 0.01, t = -0.62, P = 0.54$) or arousal PEs ($\beta = 0.02 \pm 0.01, t = 1.66, P = 0.10$). These results are presented in Supplementary Figure 6 (Fig. S6).

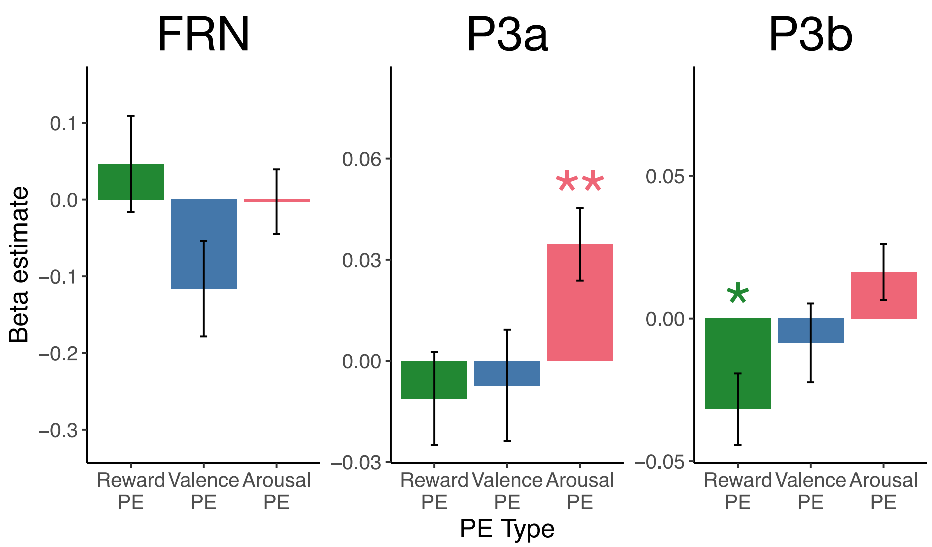


***Figure S6. Signed PEs predicting ERPs.*** *The data on each graph reflect the beta coefficient from LMMs modeling the marginal contributions of the signed value of each PE type on separate ERPs: the FRN, P3a, and P3b. Error bars reflect ±1 S.E. ***P<.001, **P<.01, *P<.05.*

*The P3a is predicted by offer extremity.* The P3a, also referred to as the novelty P3, is associated with the processing of novel or unexpected stimuli[7]. We explored the possibility that the P3a may have associations with elements of the task other than the emotional impact or reward prediction errors. Specifically, we thought that assessing "offer extremity", which measures how significantly better or worse an offer is compared to the average offer, could represent novel or unexpected information that the P3a is sensitive to. To test this, we employed a linear mixed-effects model to predict P3a amplitudes based on the linear and quadratic relationship of the offer. Prior to modeling, we standardized the offer, setting the average offer as 0 and representing deviations from the average offer as negative and positive values. The results reveal a strong prediction of P3a amplitudes by the quadratic component of the offer ($\beta = 24.33 \pm 4.63, t = 5.26, P < 0.001$), while the linear component showed no significant effect ($\beta = 3.14 \pm 4.75, t = 0.66, P = 0.51$; Fig. S7).

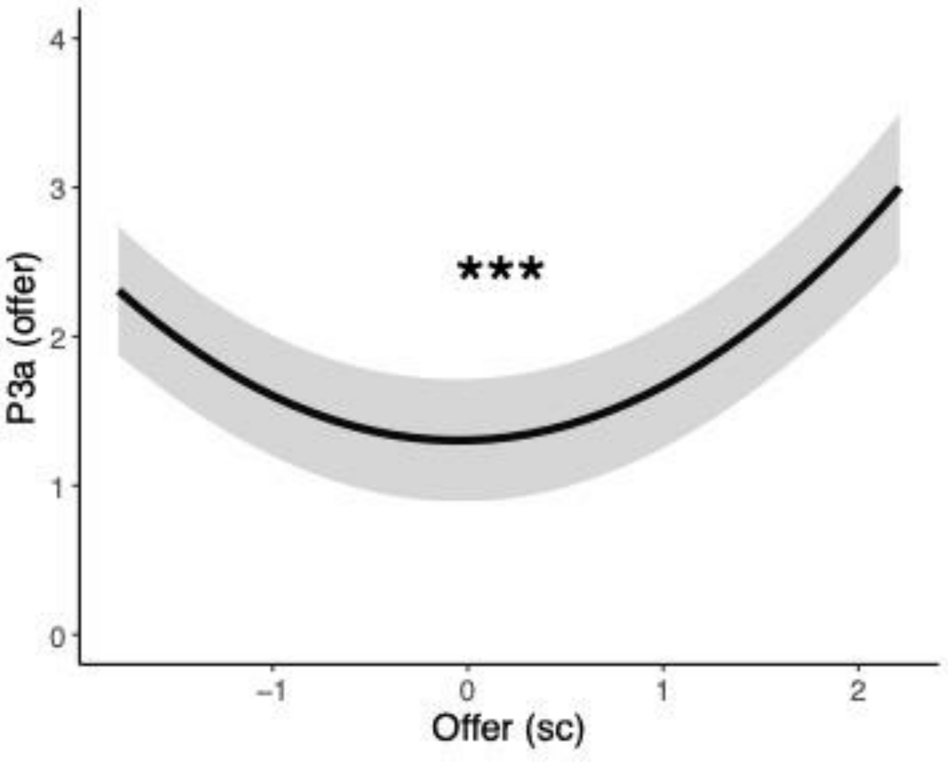


***Figure S7. P3a is sensitive to offer extremity.*** *The data reflect the predicted P3a amplitude by the linear and quadratic components of the standardized offer. Error bars reflect ±1 S.E. ***P<.001, **P<.01, *P<.05.*

*Mass univariate analysis shows separate clusters for reward and emotion PEs.* Although the ERP analysis from the manuscript offers robust evidence for a neural dissociation between emotion and reward PEs, there is a possibility that predefined ERPs might overlook crucial neural signals tied to feedback processing. To address this, we employed a data-driven method[8] that lets us identify potential electrophysiological signatures of reward, valence, and arousal PEs without the need to rely on predefined signals. We regressed each participant's offer-locked EEG signal (at every timepoint and channel) against separate variables reflecting the standardized reward, valence, and arousal PEs values, allowing us to determine the unique contribution of each PE. The beta coefficient for each PE, across every electrode (64) and timepoint (500) between -200ms and 800ms with a step size of 2ms, was calculated for each participant and then aggregated into a t-statistic representing the predictor variable's strength for the entire group. Spatiotemporal clusters were formed using a cluster-forming threshold at a p-value of $p<.01$ and the mass of each cluster was computed as the sum of the absolute t-statistics within the cluster. To control for multiple comparisons, we permuted subject betas via sign flipping to create 1000 permutations of our original data, their corresponding t-maps, and spatiotemporal clusters. A null distribution for cluster mass was created by selecting the largest cluster mass value from each permutation. Cluster significance was assessed by examination of actual cluster mass values relative to this null distribution – clusters in the $97.5^{th}$ percentile were considered significant, essentially implementing a two-tailed test.

This analysis revealed two distinct clusters for reward and valence PEs (Fig. S8A and Fig. S8B), with no significant clusters for arousal PEs. Valence PEs were represented by a positive cluster spanning 460 to 670ms after offer onset, with the most pronounced signal in the parietal area. This cluster aligns with the timing and direction of the P3b response, reinforcing the evidence that the P3b is the neural representation of valence PEs. Average feedback-locked event related potentials in the P3 electrode reveal a positive deflection for positive and negative valence PEs (Fig. S8D). Conversely, reward PEs were linked to a negative cluster ranging from 326 to 790ms, with the most robust signal in the frontal areas, including an EEG signal consistent with the timing, direction, and topography of the FRN response as shown by the average waveform in the Fz electrode (Fig. S8C). Collectively, these results strengthen the conclusion that reward and valence PEs are separately encoded in the brain.

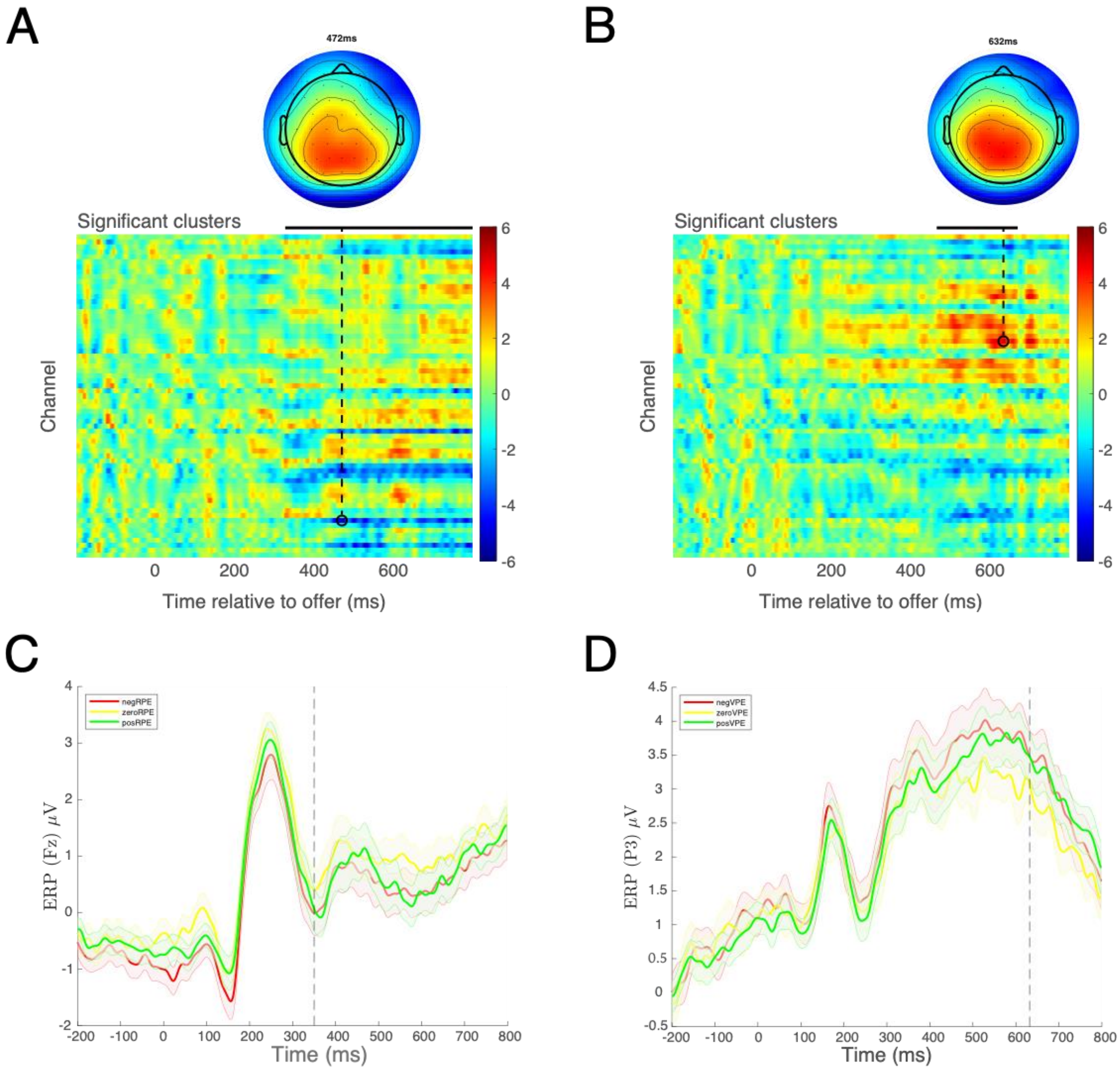


***Figure S8. Separate spatiotemporal clusters for reward and valence PEs.*** *The data on the top two graph reflect the spatiotemporal clusters that survived multiple comparisons correction as a reward PE heat plot (**A**) and a valence PE heat plot (**B**). The time and channel corresponding to the maximum absolute t-statistic for each cluster are depicted with a black circle and visualized using a topographical plot. The data on the bottom two graphs reflect the amplitudes of the Fz (**C**) and P3 (**D**) electrodes and the colors reflect bins of negative (less than -.2), zero (between -.2 and .2), and positive (greater than .2) reward (**C**) and valence (**D**) PEs. The time of the peak cluster expression for each electrode is plotted in the dashed line.*
